# The response of *Pseudomonas putida* to a complex lignolysate

**DOI:** 10.1101/764399

**Authors:** Mee-Rye Park, Yan Chen, Mitchell Thompson, Veronica T. Benites, Bonnie Fong, Christopher J. Petzold, Edward E. K. Baidoo, John M. Gladden, Paul D. Adams, Jay D. Keasling, Blake A. Simmons, Steven W. Singer

**Author notes:** Corresponding author: Joint BioEnergy Institute, 5885 Hollis Street Emeryville, CA 94608.

## Abstract

There is strong interest in the valorization of lignin derived from plant biomass to produce valuable products; however, the structural complexity of this biopolymer has been a major bottleneck to conversion. Chemical pretreatment liberates soluble fractions of lignin that may be upgraded by biological conversion. Here, ionic liquid pretreatment was employed to obtain soluble aromatic-rich fractions from sorghum, which were converted by *Pseudomonas putida* KT2440, a promising host for bioconversion of aromatics derived from lignin. Growth studies and mutational analysis demonstrated that *P. putida* growth on these soluble lignin-derived fractions, referred to as lignolysate, was dependent on aromatic monomers derived from lignin (*p*-coumarate and ferulate), but other, unknown factors in the lignolysate contributed to growth. Proteomic and metabolomic analyses provided evidence that these unknown factors were amino acids and residual ionic liquid. Proteomic measurements indicated a coordinated response in which these substrates were catabolized simultaneously. A cholinium catabolic pathway was identified and deletion of five genes in the pathway abrogated the ability of *P. putida* to grow on cholinium as a sole carbon source. This work demonstrates that lignolysates obtained through biomass pretreatment contain multiple substrates and conversion strategies for lignin-derived should take this complexity into account.

**Importance:** Lignin is one of the most abundant biopolymers on Earth and is generated as a co-product in the processing of lignocellulosic biomass. Valorization of these residual lignin streams is a promising method to enhance the economic viability of modern lignocellulosic biorefineries. In this study we developed a process to couple chemical depolymerization of lignin and biological conversion using *Pseudomonas putida* KT2440. Water-soluble and bioavailable lignolysate was obtained from sorghum and further characterized as a growth substrate for *P. putida*. Proteomic and metabolomic analyses demonstrated that P. putida metabolized other components of the lignolysate beyond monoaromatic compounds, which illuminates how microbes can process complex lignolysates obtained from plants. Understanding the underlying microbial responses in lignolysates will enable the design of rational strategies for lignin valorization.

## Introduction

Lignocellulosic biomass, which is primarily composed of cellulose, hemicellulose and lignin, represents a primary renewable feedstock for biofuel and biochemical production (1). For decades, conversion strategies have focused on the polysaccharides cellulose and hemicellulose, whereas lignin, which makes up around 15-30 wt % of biomass, is usually combusted to provide heat and electricity to support pulping operations or recovered as kraft lignin, vanillin or lignosulfonates (2, 3). Despite the fact that lignin is the only large-volume renewable aromatic feedstock on the Earth (4, 5), the bioprocessing of lignin into bioproducts is a major bottleneck because of its intrinsic heterogeneity and recalcitrance to depolymerization (5). However, with the emergence of lignocellulosic biorefineries, lignin conversion is a crucial component of integrated biorefineries with respect to economics and sustainability (5, 6).

A promising lignin valorizing strategy couples chemical lignin depolymerization with microbial catabolism of aromatic monomers by hosts that have been engineered to upgrade the depolymerized lignin (7–9), since only these low molecular weight products of lignin can be assimilated as carbon sources by microbes (9–12). To achieve this objective, chemical lignin depolymerization methods have been developed (12–15). Among these methods, a base-catalyzed depolymerization (BCD) process has been demonstrated to produce high yields of aromatic monomers, primarily *p*-coumarate and ferulate, that acylate lignin (12, 16, 17). Previously, the BCD process was employed on solid lignin-rich residue derived from corn stover via deacetylation, mechanical refining, and enzymatic hydrolysis to release lignin-derived aromatic monomers that can be further upgraded into value-added molecules (12). Ionic liquids (ILs) have been proposed as pretreatment chemicals for lignocellulosic biomass fractionation due to their highly tunable physicochemical and compatibility with biology (18, 19). Significant advances have also been made in improvement of ILs recovery and recycling to overcome high cost of ILs (18–21). Combining IL pretreatment and the BCD process may increase the bioavailable depolymerized lignin that can be converted by microbes.

*Pseudomonas putida* KT2440 is as a promising host for engineering lignin bioconversion, due to its ability to catabolize numerous aromatic compounds and its amenability to genetic manipulation. Bioconversion studies with aromatics have mostly focused on purified model compounds (7, 22), whereas the molecular mechanism of bioconversion using aromatics directly derived from plant lignin is less well-understood. Therefore, a fundamental understanding of the biological conversion of complex mixtures derived from lignin, referred to as lignolysates, is critical for rational strain engineering and upgrading of lignin-derived substrates to bioproducts. While genomic and transcriptomic analyses have been carried out to characterize the lignin-degrading mechanisms (23–25), proteomic analysis is able to offer insights into the protein abundance which is important for understanding microbial functions. Therefore, proteomic analysis is crucial for understanding the microbial conversion of lignin-derived substrates in greater depth.

In this work, we obtained size-defined soluble aromatic-rich streams from sorghum using biologically-derived ILs and assessed their biocompatibility with *P. putida*. Growth studies, mutational analyses and proteomic measurements demonstrated that the lignolysate derived from sorghum was a complex mixture composed of monoaromatics, amino acids and residual ILs that supported growth in *P. putida*.

## RESULTS

### Generation of size-defined aromatic fractions from sorghum lignin

Multiple solubilized fractions of sorghum-derived lignin were generated to identify a fraction that would be bioavailable for conversion by *P. putida* KT2440 (Figure 1). In the first approach, the soluble fraction from pretreatment with a cholinium IL, cholinium aspartate, was treated with acid and then equilibrated for 2 days at 5 °C. This acid treatment produced a precipitated solid fraction enriched in lignin (54.2%) and hemicellulose (19.2%) (Table 1). The hemicellulose was removed by enzymatic hydrolysis, providing a fraction, referred to as acid-precipitated lignin (AP lignin). Compositional analysis revealed that AP lignin, which was soluble in aqueous solution at neutral pH, was ∼70% lignin, and <5% residual carbohydrate (Table 1). Gel-Permeation Chromatography (GPC) analysis demonstrated that AP lignin contained aromatic molecules with a molecular weight distribution between 1000 and 10000 Da (Figure 2A). In a second procedure, the solid remaining after cholinium ionic liquid pretreatment was enzymatically hydrolyzed and the residual material was treated with NaOH for base catalyzed depolymerization (BCD). The aqueous fraction, referred to as base-catalyzed depolymerized lignin (BCD liquor) had 20.5% lignin and 1.5% carbohydrate (Table 1). The low recovery of lignin in the BCD liquor indicated that the majority of the lignin was depolymerized. GPC analysis confirmed that the BCD fraction had peaks corresponding to monoaromatics (0.1-0.3 kDa) as well as a broad distribution of higher molecular weight species (Figure 2(A)). The composition and concentration of the monoaromatics identified in the GPC chromatogram were further identified and quantified by LC-MS. The most abundant monoaromatic was *p*-coumarate (*p*CA) (∼1.7 g/L), and ferulate (FA) was the second most abundant monoaromatic (∼0.2 g/L) (Figure 2(B)). Other monoaromatics in the BCD liquor were syringate, *p*-hydroxybenzaldehyde, benzoate, *p*-hydroxybenzoate, vanillate, and vanillin, all at concentrations <0.1 g/L (Figure 2(C)). In contrast, AP lignin consisted of a limited number of monoaromatic, all at concentrations around <0.02 g/L (Figure 2(B) and 2(C)).

**Figure 1.**
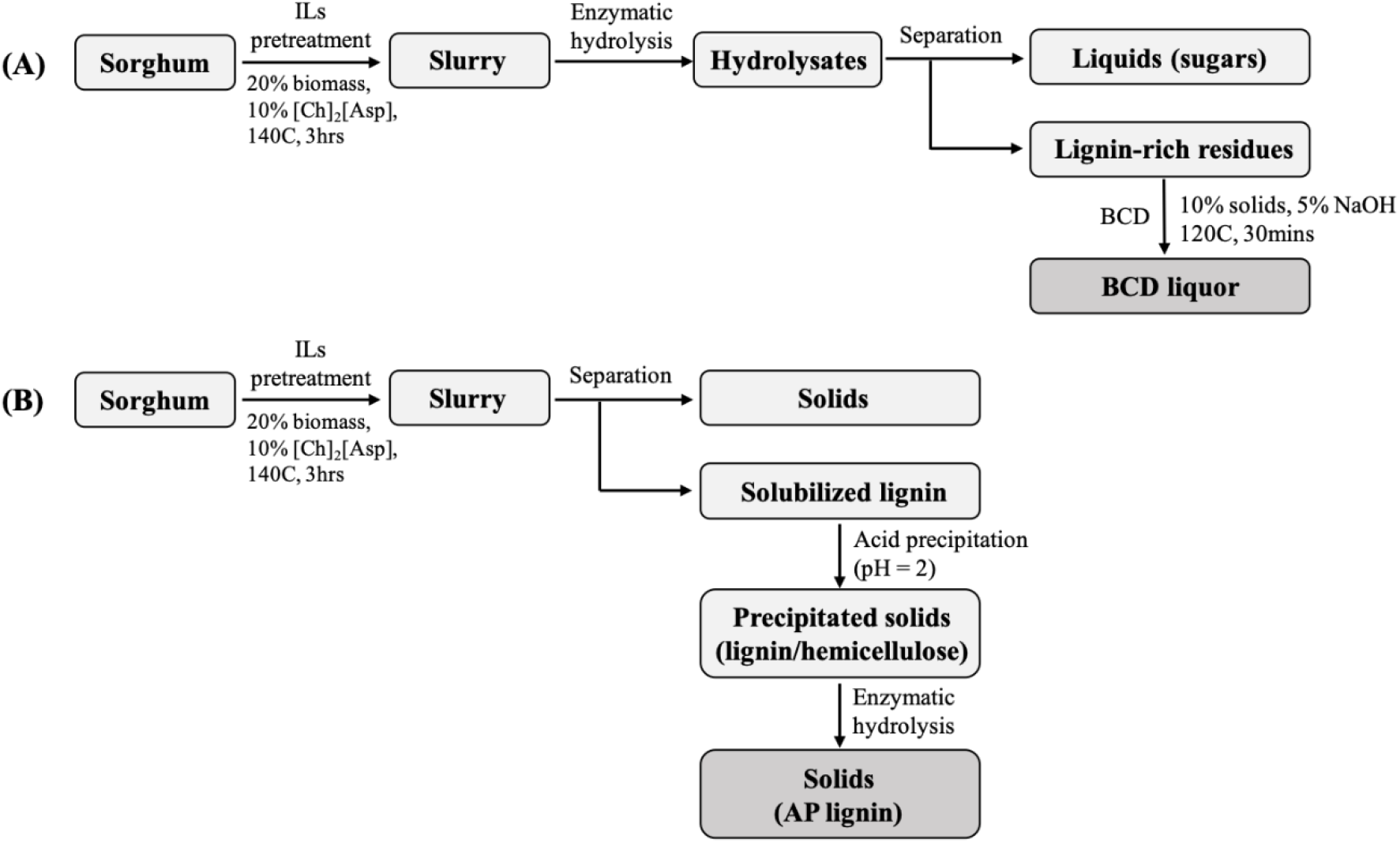
Process flow for obtaining the (A) BCD liquor and (B) AP lignin.

**Figure 2.**
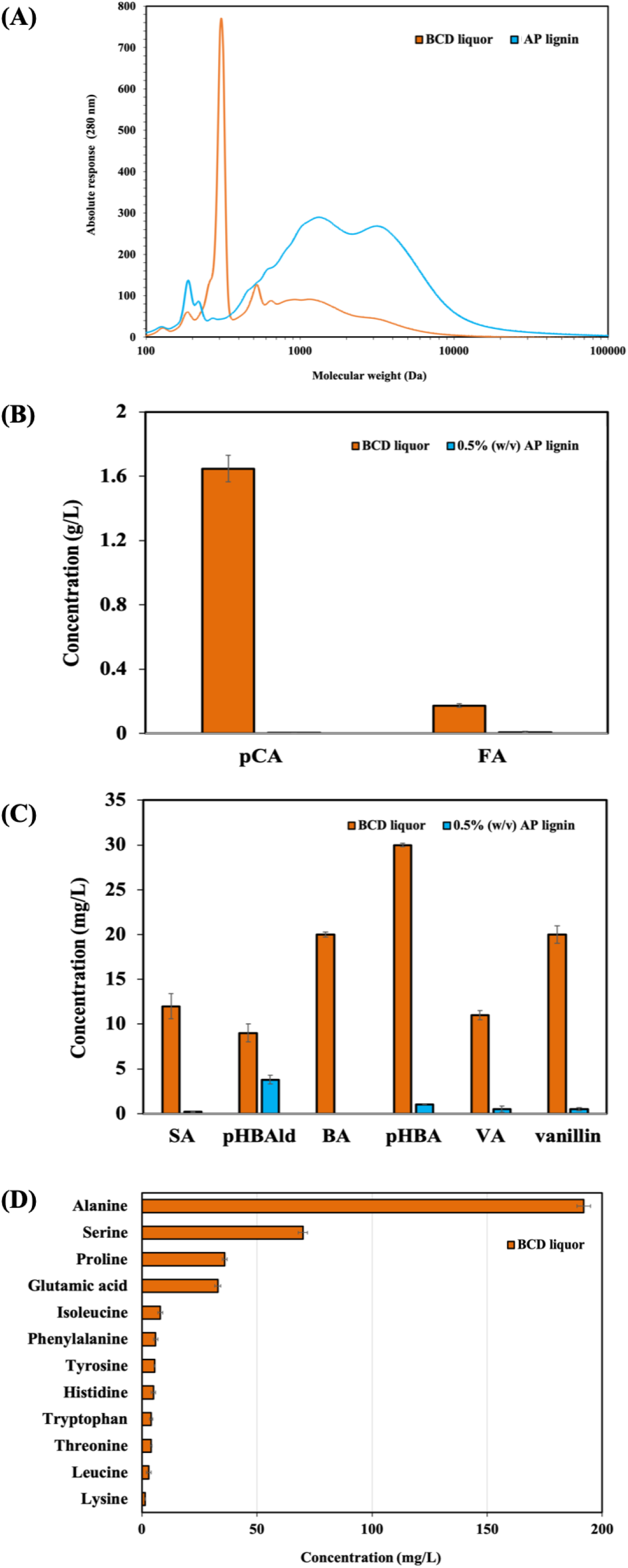
(A) Lignin molecular weight profiles (B) major lignin-derived monoaromatic compounds (C) minor monoaromatic compounds and (D) amino acid components and concentrations. *p*CA = *p*-coumarate; FA = ferulate; SA = syringate; pHBAld = *p*-hydroxybenzaldehyde; BA = benzoate; pHBA = *p*-hydroxybenzoate; VA = vanillate; VN = vanillin. The error bars indicate the error range from two technical replicates.

**Table 1.** Compositional analysis of major components in the sorghum, acid precipitated solids, acid precipitated lignin and BCD liquor (wt %).

|  | Sorghum | Precipitated solids | AP lignin | BCD liquor<sup>a</sup> |
| --- | --- | --- | --- | --- |
| Total lignin | 18.9 | 54.2 | 69.5 | 20.5 |
| Glucan | 31.5 | 1.4 | 0.5 | 0.3 |
| Xylan | 19.9 | 19.2 | 4.0 | 1.2 |
| Ash | 3 | 1 | 1.6 | 1 |
| Total | 73.3 | 75.8 | 75.6 | 23.5 |

<sup>a</sup>The composition corresponds to the soluble liquor produced at 120°C and 5% NaOH after neutralizing at pH 7 for the bacterial cultures.

### 2D HSQC NMR analysis of BCD and AP lignin

The aromatic/unsaturated (*δ*_H_/*δ*_C_ 6.0 - 8.0/90 - 160) and aliphatic (*δ*_H_/*δ*_C_ 2.5 - 6.0/50 - 90) regions of HSQC NMR spectra of the AP lignin and BCD liquor were analyzed to provide chemical information related to their composition characterized by interunit linkages (Figure 3). Signals from the aromatic ring correlations from syringyl (S) lignin (derived from sinapyl alcohol), guaiacyl (G) units (derived from coniferyl alcohol), and *p*-hydroxyphenyl (H) lignin (derived from *p*-coumaryl alcohol) were observed in the spectra of both the BCD and AP fractions. The aromatic region of the HSQC spectrum indicated that BCD fraction consisted of S (8.2%), G (88.3%), and H (3.5%) units, which represents a S/G ratio of 0.1 (Figure 3A). The AP fraction consisted of S (19%), G (73.7%), and H (7.3%) units, representing a S/G ratio of 0.26 (Figure 3B). Prominent signals corresponding to *p*CA were also observed in both the BCD and AP lignin (26, 27); in addition, signals for FA were observed in the spectrum of the AP lignin. Since the LC-MS measurements only detected free *p*CA and FA in the BCD liquor, the HSQC spectra corresponding to *p*-coumarate and ferulate found in AP lignin are likely to be *p*CA- and FA-end groups attached to oligomers. The appearance of the HSQC signals in the BCD liquor was similar to what is observed in intact lignin in the plant cell wall, whereas the AP lignin spectrum had signals corresponding to condensed S_2,6_ and condensed G_2_ units, suggesting that repolymerization reactions occurred during the process. The aliphatic/side-chain region provided important information about the lignin interunit linkages. All HSQC spectra of BCD lignin showed correlations corresponding to all of the sidechain C/H pairs for *β*-ether (*β*-*O*-4′, substructure A) and methoxyl groups, while AP lignin showed correlations corresponding to the sidechain C/H pairs for *β*-ether (*β*-*O*-4′, substructure A), resinol (*β-β*′, substructure C) units, cinnamyl alcohol (I) end groups and methoxyl groups.

**Figure 3.**
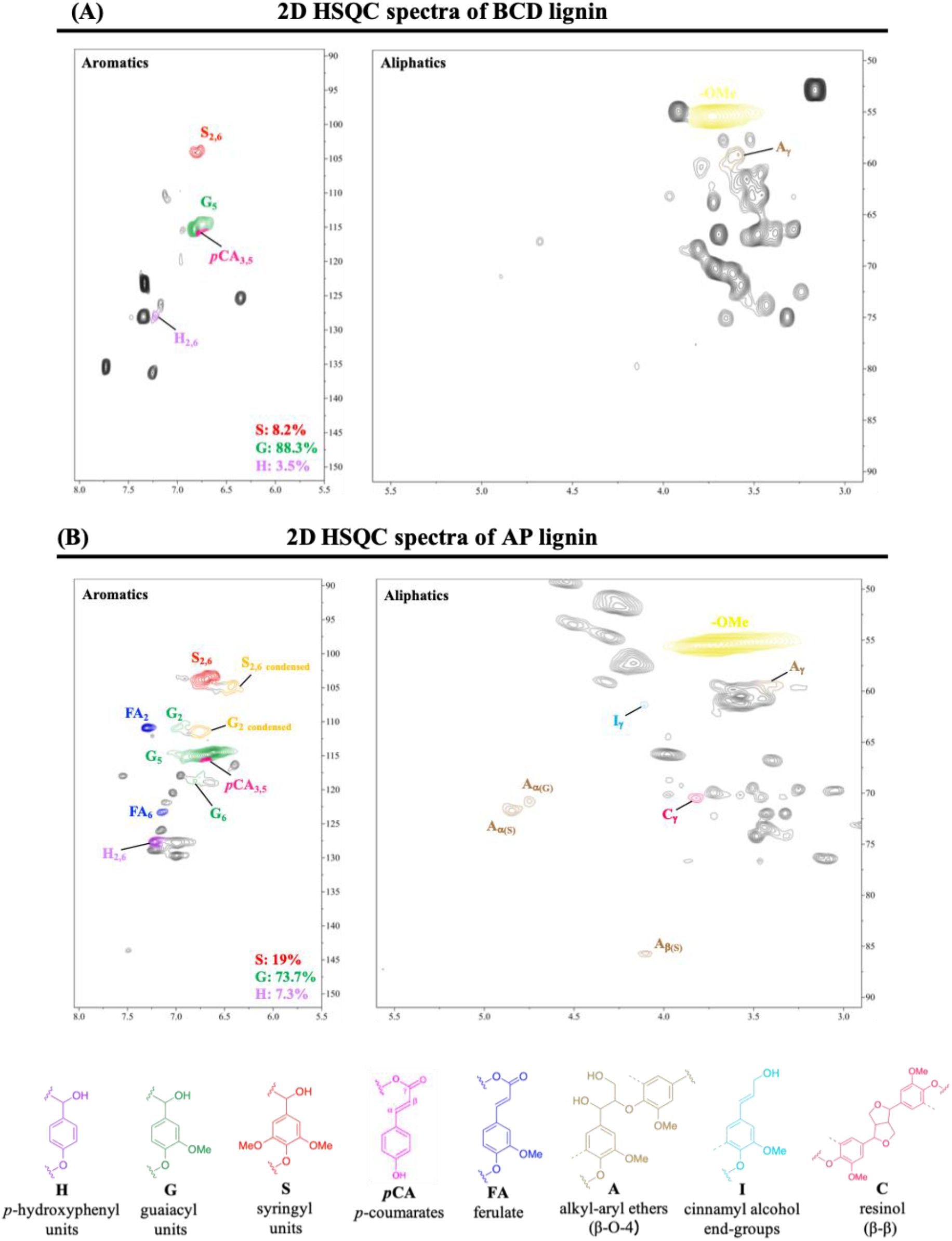
2D HSQC NMR spectra of (A) BCD liquor and (B) AP lignin. H, p-hydroxyphenyl units; G, guaiacyl units; S, syringyl units; *p*CA, *p*-coumarates; FA, ferulates; A, *β*-O-4′ alkyl-aryl ethers; I, hydroxycinnamyl alcohol endgroups; C, resinols (*β*-*β*’).

### *Pseudomonas putida* KT2440 growth on plant-derived lignolysate

Preliminary tests indicated that *Pseudomonas putida* KT2440 was capable of growth on the BCD liquor while it was incapable of growing on the AP lignin (data not shown). Bacterial growth, aromatic utilization, and lignin MW distribution before and after cultivation were investigated with 80% (v/v) BCD liquor in M9 minimal medium. The growth of *P. putida* on BCD liquor was compared to growth on a control with *p*CA as the sole carbon substrate.

Growth of *P. putida* KT2440 on BCD liquor correlated with the rapid depletion of *p*CA, FA and other monoaromatic monomers (Figure 4(A) and Figure S1). During the consumption of *p*CA and FA, transient accumulation of *p*-hydroxybenzoate and vanillate was observed as intermediates in *p*CA and FA catabolic pathways, respectively, followed by complete consumption (Figure S1). Furthermore, the MW distributions after the bacterial treatment were also performed to examine catabolism of LMW lignin derived compounds as well as depolymerization of HMW lignin. The MW profile in the uninoculated control exhibited a major peak in the LMW regions (Figure 4(B)). After microbial cultivation of the BCD liquor, the LMW species were absent in the GPC trace, consistent with the complete utilization of the aromatic monomers; however, the peaks in the HMW regions were unchanged after microbial culturing. These findings indicated that *P. putida* was able to degrade lignin-derived monoaromatic monomers but not higher molecular weight aromatics derived from lignin depolymerization. In comparison to growth on BCD liquor, *P. putida* growth on *p*CA under the same conditions and at the same concentration as present in the BCD liquor was lower (Figure 4(A)). This difference was unexpected, as it was assumed that *p*CA was responsible for almost all the *P. putida* growth observed in the BCD liquor. To determine if additional substrates were present in the BCD liquor, the ability of *P. putida* to grow on pCA and FA was abrogated by disrupting the hydroxycinnamoyl-CoA hydratase-lyase (*ech*, PP_3358) gene, whose gene product dehydrates and liberates acetyl-CoA from hydroxycinnamic acids. As expected, the ΔPP3358 mutant was not able to grow with *p*CA-as the sole carbon source (Figure 4(C)), and a major peak corresponding to LMW aromatics were still observed in the BCD liquor after microbial conversion using the *P. putida* mutant (Figure 4(D)). In parallel with the GPC profile, HPLC demonstrated that *p*CA and FA in the BCD liquor were not consumed during mutant cultivation (Figure S2). Nonetheless, the ΔPP3358 mutant strain was still capable of growing in the BCD liquor to an optical density approximately half of what was observed with the wild type *P. putida*, confirming that the BCD liquor contained additional substrates for *P. putida*.

**Figure 4.**
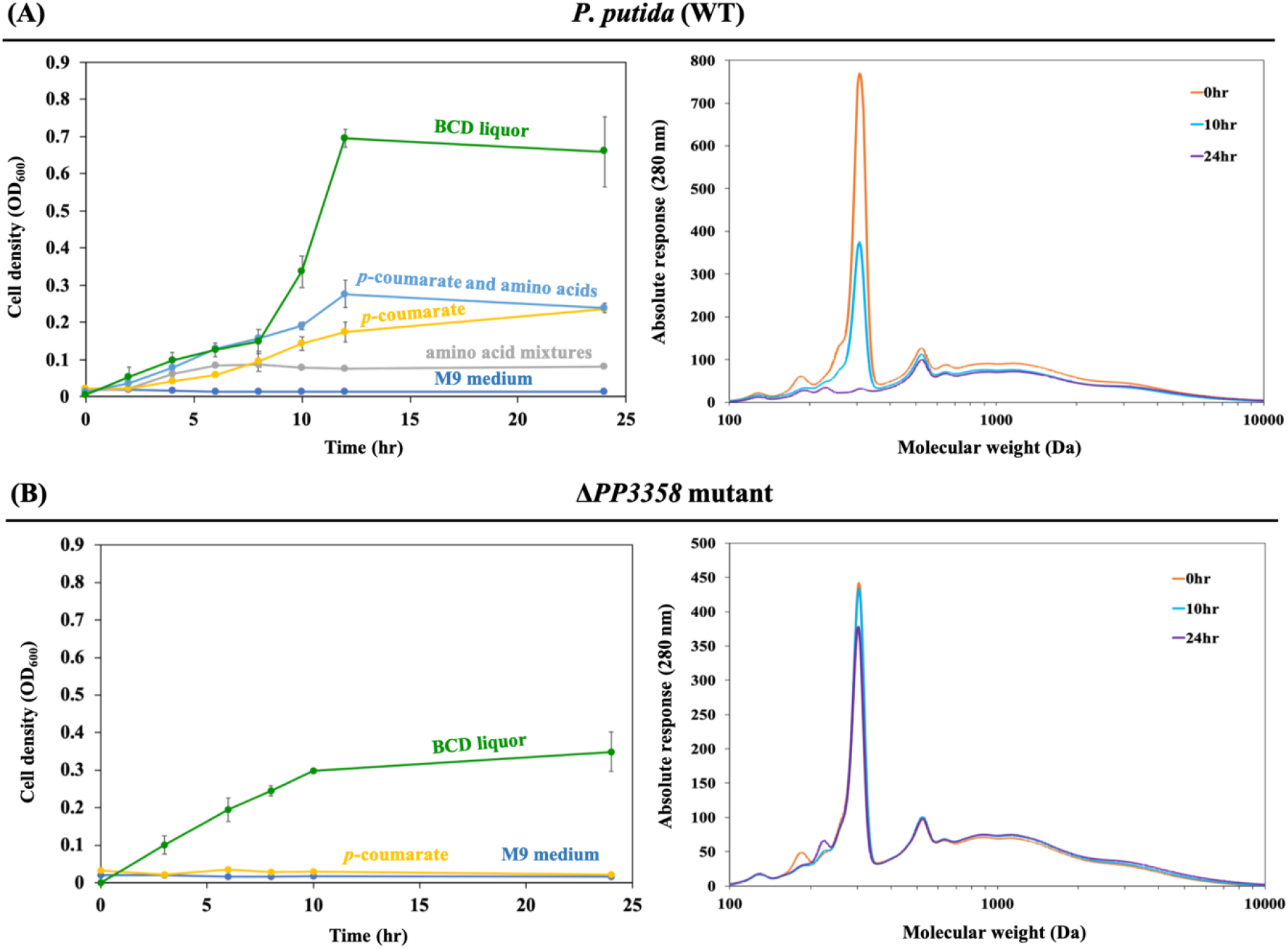
Growth profiles of *P. putida* grown in 80% (v/v) BCD liquor. Molecular weight profiles at 260 nm in BCD liquor before and after the microbial cultivation. (A) Growth profiles and (B) GPC chromatograms from BCD liquor cultivated in wt. *P. putida.* (C) Growth profiles and (D) GPC chromatograms from BCD liquor cultivated in of ΔPP3358 mutant.

A previous study demonstrated that the plant-derived amino acids in biomass hydrolysates enhanced *P. putida* growth and production of fatty acid-derived molecules (28). Therefore, amino acids liberated by the BCD process may serve as additional substrates for *P. putida*. The concentration of amino acids in the BCD liquor was measured at 0.37 g/L by LC-MS. Alanine (0.19 g/L) and serine (0.07 g/L) were the most abundant amino acids, and other low-abundance amino acids were present at ∼ 0.1 g/L in total (Figure 2(D)). A synthetic mixture mimicking the amino acids in the BCD liquor was formulated and growth of *P. putida* was tested with these amino acids. This culture (AA-only) demonstrated modest growth as the sole carbon source for growth but boosted growth when added to a medium with *p*CA (*p*CA-AA) (Figure 4(A)).

### Overview of the proteomic analysis

Since the combination of *p*-CA and the amino acids only partially recovered the growth of *P. putida* observed with the BCD liquor, global proteomic analysis of the proteins produced by *P. putida* KT2440 during growth on BCD liquor was initiated to explore the underlying microbial responses and identify determinants of increased growth with the BCD liquor as substrate. Glucose-grown cells were used as control for comparative proteomic analysis. Among 504 proteins identified, 71, 56, 34 and 39 proteins were significantly increased (log_2_ FC > 1, *p* < 0.01) in BCD liquor-, AA-, *p*CA-, and *p*CA/AA-grown cells, respectively. All of these increased proteins were clustered into 18 functional groups indicating particular metabolic processes responsible for the utilization of the different substrate sources (Figure S3). According to the COG analysis, significantly increased proteins in *P. putida* in response to BCD liquor were grouped into the categories “energy production and conversion”, “lipid transport and metabolism”, “amino acid transport and metabolism” and “secondary metabolites biosynthesis, transport and catabolism” in cells grown in the BCD liquor.

### Differentially increased proteins in *P. putida* KT2440 grown in BCD liquor

Growth of *P. putida* in a M9 minimal medium containing BCD liquor led to the significant induction of proteins associated with aromatic catabolic and *β*-ketoadipate pathways (Figure 5 and Table S1). More specifically, the proteins involved in the conversion of *p*-coumarate and ferulate to hydroxybenzoate: 4-coumarate: CoA ligase (Fcs), hydroxycinnamoyl-CoA hydratase-lyase (Ech) and vanillin dehydrogenase (Vdh) were significantly increased (5.12- to 7.51-log_2_ FC) when cells were grown in BCD liquor compared to the control culture grown from sugar only. 4-hydroxybenzoate hydroxylase (PobA), which transforms 4-hydroxybenzoate into protocatechuate, was also significantly increased (6.75-log_2_ FC). The subsequent enzymes encoded by the *pca* genes further catalyze the protocatechuate ortho-cleavage pathway (29). The *pca* genes are arranged in four different clusters, *pcaHG*, *pcaBDC*, *pcaIJ*, and *pcaF*. Herein, enzymes (PcaHG, PcaB and PcaD) belonging to the protocatechuate branch of the *β*-ketoadipate pathway were significantly increased (1.41- to 5.96-log_2_ FC). 4-carboxymuconolactone decarboxylase (PcaC) required for the transformation of 4-carboxymucono-lactone to beta-ketoadipate-enol-lactone was not detected in this study. Lastly, proteins (PcaIJ and PcaF) involved in two further steps of converting *β*-ketoadipate into tricarboxylic acids (TCA) cycle intermediates were also significantly increased (3.04 and 2.88-log_2_ FC, respectively). Similar changes in the level of these enzymes involved in the aromatic catabolic pathway was also observed in *p*CA-only (3.3- to 7.71-log_2_ FC) and *p*CA/AA (1.89- to 7.84-log_2_ FC) controls (Figure S4(B),(C) and Table S2). Degradation of *p*-coumarate through the protocatechuate-branch of the *β*-ketoadipate pathway yields acetyl-CoA and succinyl-CoA, which enter the TCA cycle (29). TCA cycle enzymes citrate synthase (GltA, 0.76-log_2_ FC), aconitate hydratase (AcnA-2, 2.07-log_2_ FC), isocitrate dehydrogenase (Icd, 0.96-log_2_ FC), succinate dehydrogenase (SdhAB, 0.93- to 1.16-log_2_ FC) and the first enzyme of glyoxylate shunt (isocitrate lyase (AceA), 4.51-log_2_ FC) were at significantly higher abundance when cells were grown in BCD liquor. Similar results were observed in succinate dehydrogenase and glyoxylate shunt enzymes in *p*CA-only (1.15- to 4.4-log_2_ FC) and *p*CA/AA (0.6- to 3.99-log_2_ FC) controls (Figure S4(B),(C) and Table S2).

**Figure 5.**
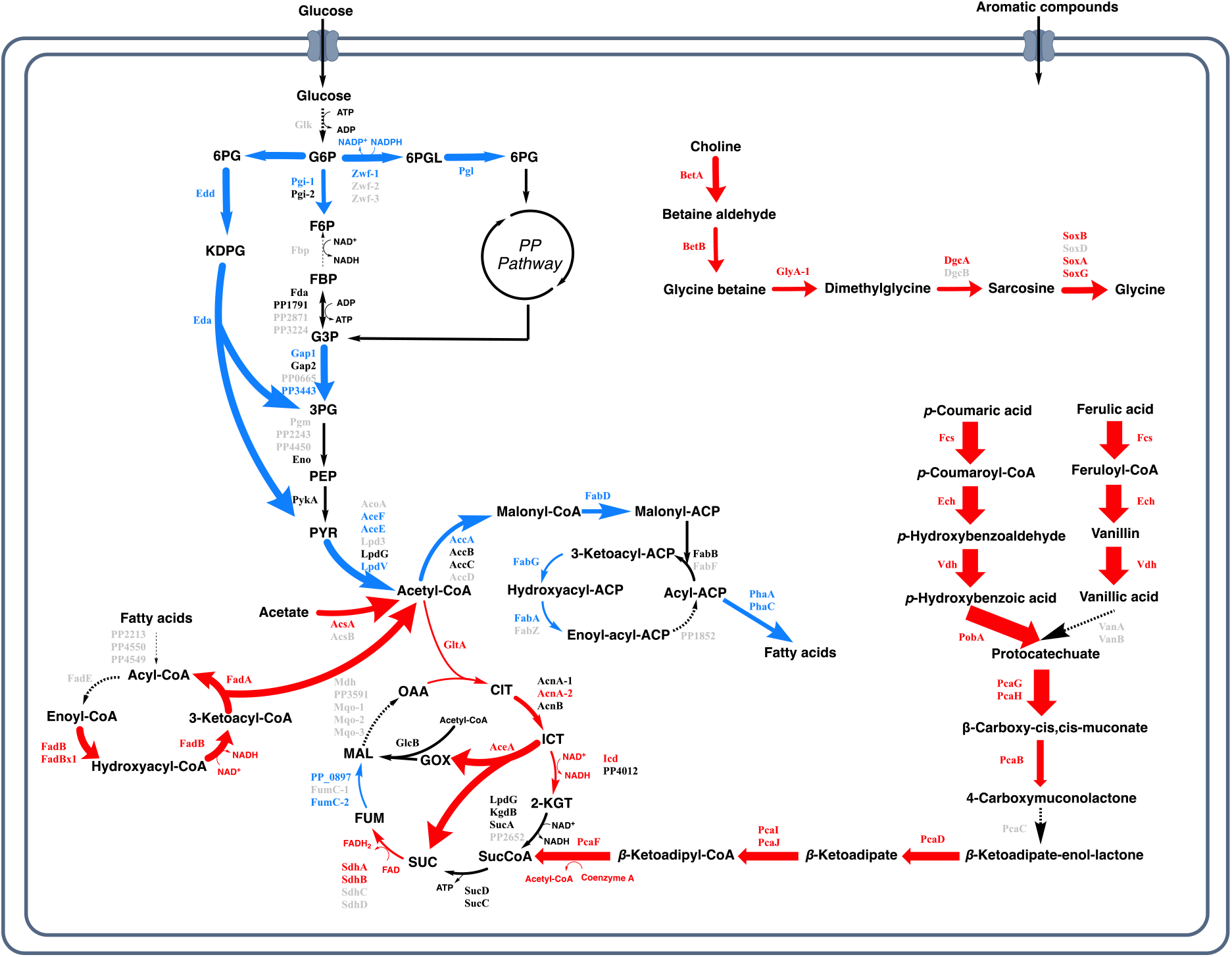
Differentially abundance of *P. putida* proteins grown in the presence of BCD liquor compared to a glucose control. Increased fold changes of proteins/pathways are marked in red, and decreased fold changes of proteins/pathways are marked in Blue. The abundance of proteins/pathways indicated in black were not significantly changed in the BCD liquor. Proteins/pathways indicated in gray and dashed lines were not observed in our proteomics survey. The abbreviations used are: ED pathway, Entner-Doudoroff pathway; EMP pathway, Embden-Meyerhof-Parnas pathway; PP pathway, pentose phosphate pathway; G6P, glucose-6-P; 6PG, 6-phosphogluconate; 6PGL, 6-P-glucono-1,5-lactone; F6P, fructose-6-P; FBP, fructose-1,6-P2; G3P, glyceraldehyde-3-P; 3PG, glycerate-3-P; PEP, phosphoenolpyruvate; Pyr, pyruvate; OAA, oxaloacetate; CIT, citrate; ICT, isocitrate, 2-KGT, 2-ketoglutarate; SucCoA, succinyl-CoA; SUC, succinate; FUM, fumarate; MAL, malate.

**Table 2.**
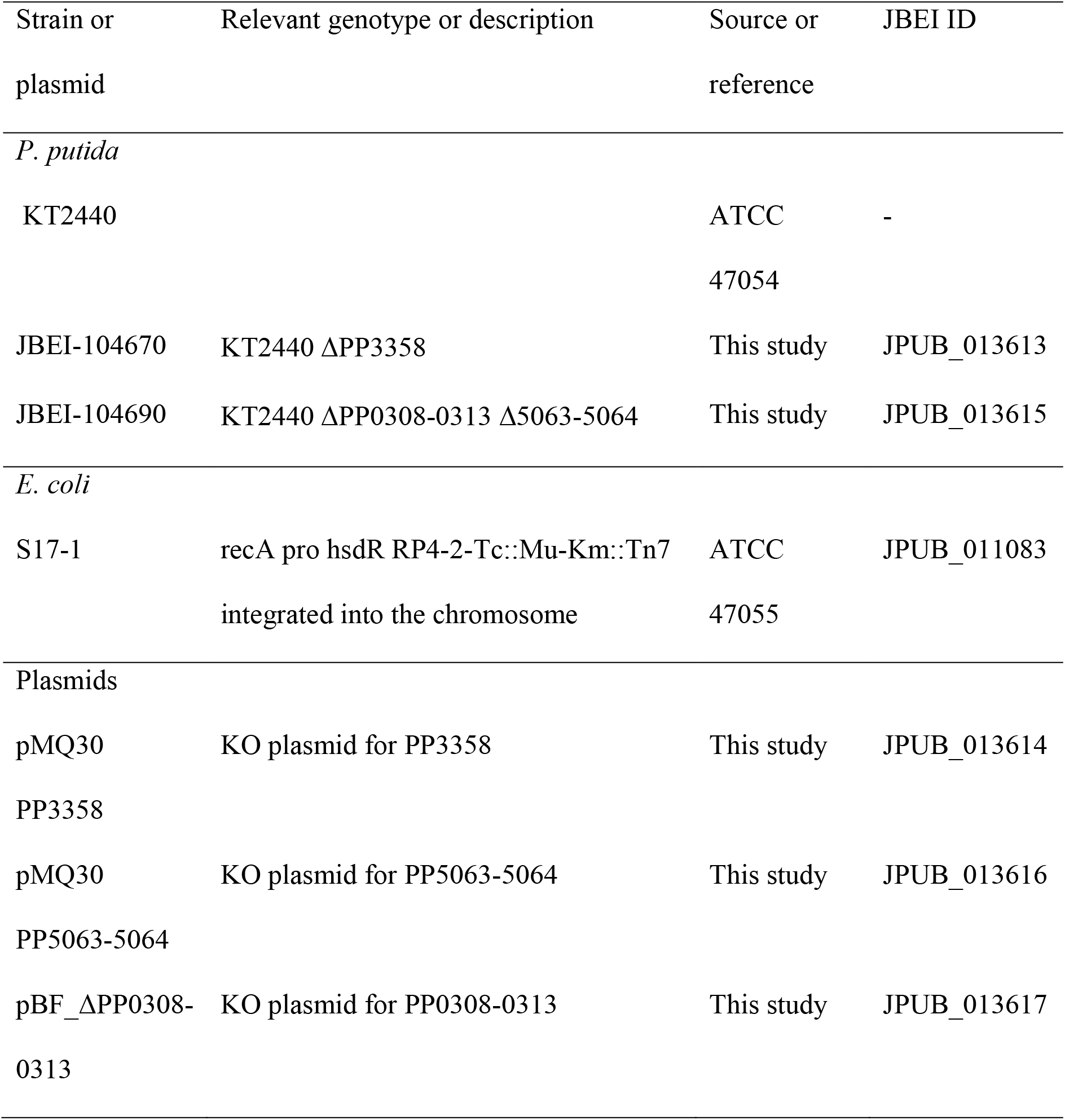
Bacterial strains and plasmids used in this study.

| Strain or<br>plasmid | Relevant genotype or description | Source or<br>reference | JBEI ID |
| --- | --- | --- | --- |
| *P. putida* |  |  |  |
| KT2440 |  | ATCC<br>47054 | - |
| JBEI-104670 | KT2440 ΔPP3358 | This study | JPUB_013613 |
| JBEI-104690 | KT2440 ΔPP0308-0313 Δ5063-5064 | This study | JPUB_013615 |
| *E. coli* |  |  |  |
| S17-1 | recA pro hsdR RP4-2-Tc::Mu-Km::Tn7<br>integrated into the chromosome | ATCC<br>47055 | JPUB_011083 |
| Plasmids |  |  |  |
| pMQ30<br>PP3358 | KO plasmid for PP3358 | This study | JPUB_013614 |
| pMQ30<br>PP5063-5064 | KO plasmid for PP5063-5064 | This study | JPUB_013616 |
| pBF_ΔPP0308-0313 | KO plasmid for PP0308-0313 | This study | JPUB_013617 |

An additional substrate that may contribute to *P. putida* growth are plant-derived fatty acids. Fatty acids are essential components of membranes and are important sources of metabolic energy, which can either be degraded via *β*-oxidation pathway or used as precursors for important building blocks such as phospholipids. GC-MS measurements demonstrated low levels of fatty acids in the BCD liquor, with octadecanoic acid being the most abundant (0.01 g/L). Despite the low abundance, proteins involved in fatty acid *β*-oxidation (FadAB) showed significant increases in abundance (1.21- to 4.33-log_2_ FC) in the proteome of the culture with the BCD liquor. However, these proteins were also increased in the proteome of the AA-only and *p*CA-AA cultures, suggesting that the increased abundance of *β*-oxidation proteins may arise as a result of amino acid metabolism. Proteins involved in amino acid transport and their metabolism were observed in higher levels compared to glucose-only control. For example, glutaminase (AnsB, 1.98-log_2_ FC), glutamate/aspartate ABC transporter-periplasmic binding protein (GltI, 2.27-log_2_ FC) and choline/betaine/carnitine ABC transporter-substrate binding protein (BetX, 1.88-log_2_ FC) were significantly increased probably due to the utilization of plant-derived amino acids. The enzymes were also increased in the control groups with amino acids present (2.06- to 2.69- log_2_ FC in AA-only; 0.56- to 2.0-log_2_ FC in *<u>p</u>*CA/AA).

Another possible substrate in the BCD liquor could be residual IL from the initial pretreatment. We considered this an unlikely possibility, since the solids remaining after both IL pretreatment and enzymatic hydrolysis was extensively washed. However, other *Pseudomonas* species have been shown to catabolize cholinium (30–32), and growth on *P. putida* on 0.2% cholinium aspartate as the sole carbon source was demonstrated (Figure S5). To determine if the catabolism of cholinium occurred during growth on the BCD liquor, a cholinium catabolic pathway was identified in the *P. putida* genome by reference to a characterized pathway in *Pseudomonas aeruginosa* (30, 31). Cholinium is oxidized to glycine betaine by genes encoding choline dehydrogenase (BetA) and betaine aldehyde dehydrogenase (BetB), followed by demethylation of glycine betaine to dimethylglycine by serine hydroxymethyltransferase (GlyA-1). The demethylation of dimethylglycine is carried out by DgcA and DgcB. Sarcosine demethylation is conducted by a heterotetrameric enzyme, SoxBDAG. Proteomic analysis revealed that the some of the proteins of the cholinium catabolic pathway was significantly increased in BCD liquor (Table S1): BetA (2.93-log_2_ FC), BetB (4.02-log_2_ FC), GlyA-1 (2.58-log_2_ FC), DgcA (2.91- log_2_ FC) and SoxABG (3.06- to 5.05-log_2_ FC). The catabolism of cholinium was also confirmed by LC-MS analysis of the BCD liquor, which indicated that ∼0.2 g/L of cholinium was present in the BCD liquor and was consumed during the course of the cultivation with the BCD liquor. The ability of *P. putida* to grow on cholinium aspartate was eliminated by deleting *dgcAB* (ΔPP_0308-0313) and *betBA* (ΔPP_5063-5064) in KT2440 (Figure S5).

## DISCUSSION

The biocompatible lignin depolymerization method described here enables a route to biologically upgrading lignin into value-added bioproducts, a process that could potentially be run in parallel with sugar conversion or as a separate stream. We employed choline-based IL pretreatment to obtain solubilized and insoluble lignin fractions that were then further processed into size-defined fractions of lignin, which were examined for microbial utilization. In one approach, a low molecular weight fraction was obtained by BCD reaction of the solid fraction after saccharification, while a relatively high molecular weight fraction was produced by another approach using acid precipitation of the IL-solubilized lignin. HSQC NMR of the AP lignin was consistent with condensation of lignin, which may arise during IL pretreatment or subsequent acid precipitation (33), whereas BCD provided depolymerized lignin streams without condensation.

The substrates *p*-coumarate and ferulate were the main lignin-derived aromatic monomers present in BCD liquor, but *p*-coumarate was present at much higher levels than ferulate. The higher abundances of *p*-coumarate compared to ferulate support previous observations that *p*-coumarate is predominantly attached to the lignin while ferulate is mostly attached to the polysaccharides. Previous studies have indicated that the bulk of *p*-coumarate is esterified to the lignin side chains and acylates the γ-OH of the lignin side chain in grasses (34–37). On the other hand, ferulate has been shown to be involved in lignin-polysaccharides linkages (38). While previous studies focused mainly only on aromatic compounds in BCD liquor (12, 13, 17), this study further revealed that BCD liquor contains fatty acid and residual ILs (choline) as well as aromatics and amino acids, which revealed not only specific, differentially increased proteins of KT2440 using the mass spectrometry-based proteomic approach, but also aromatic-independent growth in by a *P. putida* mutant strain that was unable to metabolize *p*-coumarate and ferulate. The amino acids in BCD liquor were probably liberated during the BCD reaction by hydrolysis of plant proteins in the solid fraction after one-pot pretreatment. Some covalent linkages have also been demonstrated between lignin, polysaccharides and structural proteins of grass cell walls (39). The plant-derived amino acids (serine, valine, aspartate, phenylalanine and tryptophan) were also shown in hydrolysates obtained from *Arabidopsis*, switchgrass and sorghum (28). Likewise, in this study, free fatty acids in BCD liquor were probably produced by BCD reaction since a common feature of the lipid membrane components in plant cells is esters of moderate to long chain fatty acids, and acid or base-catalyzed hydrolysis yields the component fatty acid from the lipid components (40, 41).

When *P. putida* KT2440 were grown in the BCD liquor, complete utilization of aromatic monomers was observed during aromatic catabolism corresponding with the disappearance of LMW lignin peaks as shown by GPC, which is in agreement with a previous study (12). We observed the complete list of proteins responsible for aromatic catabolism via the protocatechuate ortho-cleavage pathway and *β*-ketoadipate pathway. It should be noted that the proteins participating in the catabolic pathways for aromatic compounds to central metabolism increased significantly in the proteome, indicating that lignin-derived aromatics in the BCD liquor are the main carbon and energy sources, which is consistent with the observations in the presence of *p*-coumarate (*p*CA-only and *p*CA/AA controls). However, depolymerization of HMW lignin by *P. putida* KT2440 was not observed in our system. These results are in contrast to those of a previous study (8), which reported simultaneous depolymerization of HMW lignin and aromatic catabolism in alkaline pretreated liquor by *P. putida* KT2440. Notably, no ligninolytic enzymes were detected in the proteomes generated during cultivation on the BCD liquor.

Unexpectedly, in the BCD liquor, we observed significant upregulation of the fatty acid *β*-oxidation pathway and acetyl-CoA synthetase, both of which may enable an increase of acetyl-CoA levels and a subsequent increase in carbon flux towards the glyoxylate shunt and TCA cycle, respectively(45). Considering the elevated proteins levels involved in *β*-oxidation in the presence of amino acids (AA-only and *p*CA/AA controls), the significant increase of fatty acid *β*-oxidation pathway in BCD liquor was probably due to amino acid metabolism.

Additionally, TCA cycle dehydrogenases (Icd, SdhA and SdhB) were increased. All of the dehydrogenases mentioned above are involved in the NADH or FADH_2_ generation, and their oxidation leads to ATP production. Moreover, we observed significantly increased isocitrate lyase which is consistent with all controls. Isocitrate lyase catalyzes the formation of glyoxylate and succinate from isocitrate as a key step of the glyoxylate cycle (46). The glyoxylate shunt is known to be increased when acetyl-CoA is a product of a metabolic pathway, for example via degradation of fatty acids, acetate or alkanes (45) and bypassing a portion of the TCA cycle conserves carbon for gluconeogenesis while simultaneously diminishing the flux of electrons funneled into respiration (47). Taken together, it seems likely that *P. putida* needs to increase the availability of acetyl-CoA and/or electron carriers when growing on the BCD liquor to obtain energy and precursors for cellular biosynthesis. It is involved in the metabolic adaptation in response to environmental changes, which operates as an anaplerotic route for replenishing the TCA cycle during growth under glucose limitation (46, 48).

This study highlights the potential for biological upgrading of the soluble and biocompatible lignin-rich substrates obtained by ILs pretreatment and BCD process. Besides lignin-derived compounds, other cell wall components and residual cholinium-based ILs were further identified in the plant-derived lignolysate (BCD liquor herein), thereby a wide variety of proteins increased in *P. putida* KT2440 in response to the presence of lignolysate. These findings can provide an enhanced understanding of their metabolic pathways for carbon utilization by *P. putida*, thus forming a basis for the design of future metabolic engineering in bioconversion of lignocellulose, where multiple engineering metabolic pathways will be considered in synthetic biology methods.

## MATERIALS AND METHODS

### Materials

Sorghum was provided by Idaho National laboratory and ground using a Wiley mill through a 2mm screen and separated by a vibratory sieve system (Endecotts, Ponte Vedra, FL, USA). The commercial enzyme products Cellic® CTec3 and Cellic® HTec3 were gifts from Novozymes, North America (Franklinton, NC, USA). Cholinium aspartate was purchased from IoLiTec (Heilbronn, Germany).

### One-pot pretreatment and base-catalyzed depolymerization

IL pretreatment was conducted to process sorghum in a miniclave drive reactor (Buchiglas, Switzerland), containing 70 g wet biomass, 35 g of cholinium aspartate ([Ch]_2_[Asp]) and 245 g of DI water to give 20% (w/w) biomass loading at 140°C for 3hr. Following pretreatment, pH was adjusted to 5.5 with concentrated HCL and enzymatic hydrolysis was conducted with 50mM citrate buffer. Enzyme mixtures (Cellic® CTec3 and HTec3 at ratio of 9:1 v/v) were added to the pH-adjusted slurry. The enzymatic saccharification step was operated at 50 °C for 72 h with constant agitation on an Enviro Genie SI-1200 Rotator platform (Scientific Industries, Inc., Bohemia, NY). The hydrolysates were centrifuged at 15300xg to separate the solid and liquid phases. Solids were washed 10 times with 200 mL of DI water and lyophilized in a FreeZone Freeze Dry System (Labconco, Kansan City, MO, USA).

For the BCD reaction, the lyophilized substrate was added as 10% (w/v) solids to a 5% NaOH solution, loaded into a 350 mL stainless steel Miniclave drive 3 pressure reactor (Buchiglas, Switzerland), which was equipped with an impeller and temperature controller. The reaction proceeded through a 35 min ramp from 25 to 120 °C, a 30 min reaction at 120 °C, and a 25 min ramp from 120 to 40 °C, while keeping the stirrer speed constant at 1500 rpm as described previously (12). After the BCD reaction, the pH of the resultant liquor was adjusted to 7 with 5 N H_2_SO_4_, and the aqueous fraction was separated from the remaining solids by centrifugation. The aqueous fractions from all the BCD reactions were sterilized by surfactant-free cellulose acetate (SFCA) filtration chambers (Thermo Fisher Scientific, Waltham, MA, USA) for bacterial growth assays.

### IL pretreatment and acid precipitation

Cholinium aspartate pretreatment of sorghum was performed in a Parr reactor containing 70g wet biomass, 35 g of [Ch]_2_[Asp] and 245 g of DI water to give 20% (w/w) biomass loading at 140 °C for 3hr. After pretreatment, the slurry was transferred to 500 mL centrifuge bottles tubes (Nalgene, Rochester, NY, USA), centrifuged and filtered through 5-10 μm polypropylene bag mesh (The Western States Machine Company, Hamilton, OH, USA) to separate liquid and solid phases. The liquid phase was adjusted to pH=2 with concentrated H_2_SO_4_ and the liquids were stored at 5 °C for 48hr to obtain an acid precipitate. The mixture was transferred to 500 mL centrifuge bottles tubes and centrifuged at 15300 x *g* to obtain acid-precipitated solids. The recovered solids were further washed ten additional times with distilled water at pH 2 and the materials were lyophilized. Enzymatic hydrolysis of lyophilized solids was further carried out at 50 °C in an incubator shaking at 200 rpm for 72 h with an enzyme mixture of Cellic® CTec3 and HTec3 at ratio of 1:1 v/v, respectively, after pH adjustment (pH = 5).

### Gel-Permeation Chromatography analysis

GPC was used to determine the relative molecular weight (MW) distribution of the lignin in AP lignin and BCD liquor before and after microbial cultivation. The methodology for the GPC analysis employed in this work has been reported previously (13). Briefly, samples consisting of 20 mg of dried material from culture supernatants were acetylated with a mixture of acetic anhydride (0.5 mL) and pyridine (0.5 mL) at 40 °C for 24 h. Methanol was added (0.2 mL) to terminate the reaction, and all solvents were evaporated with nitrogen gas. The samples were dried in a vacuum oven at 40 °C overnight, dissolved in tetrahydrofuran (THF), and filtered with 0.45 μm polytetrafluoroethylene (PTFE) filters (GE Healthcare Life Sciences, USA). GPC analysis was performed using an Agilent HPLC with 3 GPC columns (Polymer Laboratories, 7.5 mm i.d. × 300 mm length) packed with polystyrene-divinylbenzene copolymer gel (10 μm beads) with THF as eluent at a flow rate of 1 mL/min at 35 °C. Absorbance at 260 nm was quantified with a diode array detector, and the retention time was converted to molecular weight using a calibration curve made with polystyrene standards.

### Compositional analysis

Total sugars, lignin extractives and ashes from untreated sorghum, precipitated solids after acid treatment, AP lignin and BCD liquor were determined according to NREL protocols (49). The samples (100 mg dry material per mL) were subjected to two-step acid hydrolysis: the first step with 72% (w/w) H_2_SO_4_ at 30 °C for 1 h and the second step conducted in the presence of 4% (w/w) sulfuric acid at 121 °C for 1 h. The amount of monomeric sugars was determined from the filtrate by high performance liquid chromatography (HPLC) as described below. The amount of glucan and xylan was calculated from the glucose and xylose content multiplied by the anhydro correction factors of 162/180 and 132/150, respectively. After hydrolysis, the hydrolysates were filtered through a filter crucible (pore size 4; Schott, Germany). The solids remaining after two stage acid hydrolysis, Klason lignin and ash, were measured gravimetrically while the acid-soluble lignin (ASL) content was quantified by measuring the UV absorbance of the acid hydrolysis supernatant at 240 nm. Ash was determined through changes in weight upon heating to 575 °C. The reported values represent the average of three technical replicates.

Monomeric sugars in the supernatant collected and quantified using an Agilent HPLC 1260 Infinity equipped with a 300 × 7.8 mm Aminex HPX 87 H column (Bio-Rad, Hercules, CA, USA) and Refractive Index Detector heated at 35 °C. An aqueous solution of H_2_SO_4_ (4 mM) was used as the mobile phase (0.6 mL min−1, column temperature 50 °C). The injection volume was 20 μL with a run time of 20 min.

### 2D ^13^C-^1^H HSQC NMR spectroscopy

Lyophilized samples were ball-milled, solubilized in 4:1 DMSO-*d_5_*/pyridine-*d_6_*, and then analyzed by two-dimensional (2D) ^1^H-^13^C heteronuclear single-quantum coherence (HSQC) nuclear magnetic resonance (NMR) spectroscopy as previously described (50). Briefly, ball-milled samples were placed in NMR tubes with 600 μL DMSO-*d_6_*/pyridine-*d_5_*. The samples were sealed and sonicated to homogeneity in a Branson 2510 table-top cleaner (Branson Ultrasonic Corporation, Danbury, CT). The temperature of the bath was closely monitored and maintained below 55 °C. HSQC spectra were acquired at 25 °C using a Bruker Avance-800 MHz instrument equipped with a 5 mm inverse gradient ^1^H/^13^C cryoprobe using the “hsqcetgpsisp2.2” pulse program (ns = 200, ds = 16, number of increments = 256, d1 = 1.0 s). Chemical shifts were referenced to the central DMSO peak (δ_C_/δ_H_ 39.5/2.5 ppm). Assignment of the HSQC spectra is described elsewhere (51, 52). Changes in lignin structural characteristics were determined based on volume integration of HSQC spectral contour correlations using the Bruker’s Topspin 3.1 processing software.

### LC-MS analysis of phenolic compounds

All metabolites were quantified using HPLC-electrospray ionization (ESI)-time-of-flight (TOF) mass spectrometry (MS). An aliquot of the culture medium was cleared by centrifugation (21,000 *x g*, 5 min, 4°C) and filtered using Amicon Ultra centrifugal filters (3,000 Da MW cut off regenerated cellulose membrane; Millipore, Billerica, MA, USA) prior to analysis. The separation of metabolites was conducted on the fermentation-monitoring HPX-87H column with 8% cross-linkage (150-mm length, 7.8-mm inside diameter, and 9-μm particle size; Bio-Rad, Richmond, CA, USA) using an Agilent Technologies 1100 Series HPLC system. A sample injection volume of 10 μl was used throughout. The sample tray and column compartment were set to 4 and 50°C, respectively. Metabolites were eluted isocratically with a mobile-phase composition of 0.1% formic acid in water at a flow rate of 0.5 ml/min. The HPLC system was coupled to an Agilent Technologies 6210 series time-of-flight mass spectrometer (LC-TOF MS) via a MassHunter workstation (Agilent Technologies, CA, USA). Drying and nebulizing gases were set to 13 liters/min and 30 lb/ in^2^, respectively, and a drying-gas temperature of 330°C was used throughout. ESI was conducted in the negative ion mode and a capillary voltage of −3,500 V was utilized. All other MS conditions were described previously (53).

### LC-MS analysis of amino acids

For the measurement of plant-derived amino acids in the BCD fraction, liquid chromatographic separation was conducted using a Kinetex HILIC column (100-mm length, 4.6-mm internal diameter, 2.6-μm particle size; Phenomenex, Torrance, CA) using a 1200 Series HPLC system (Agilent Technologies, Santa Clara, CA, USA) as described previously (28). The injection volume for each measurement was 2 μL. The sample tray and column compartment were set to 6°C and 40°C, respectively. The mobile phase was composed of 20 mM ammonium acetate in water (solvent A) and 10 mM ammonium acetate in 90% acetonitrile and 10% water (solvent B) (HPLC grade, Honeywell Burdick & Jackson, CA, USA). Ammonium acetate was prepared from a stock solution of 100 mM ammonium acetate and 0.7 % formic acid (98-100% chemical purity, from Sigma-Aldrich, St. Louis, MO, USA) in water. Amino acids were separated with the following gradient: 90% to 70%B in 4 min, held at 70%B for 1.5 min, 70% to 40%B in 0.5 min, held at 40%B for 2.5 min, 40% to 90%B in 0.5 min, held at 90%B for 2 min. The flow rate was varied as follows: held at 0.6 mL/min for 6.5 min, linearly increased from 0.6 mL/min to 1 mL/min in 0.5 min, and held at 1 mL/min for 4 min. The total run time was 11 min. The mass spectrometry parameters have been previously described (54).

### GC-MS analysis for fatty acid

Fatty acid was quantified using a method as described previously (55). Specifically, 0.5 mL of supernatant was acidified with 50 μL of concentrated HCl (12N). The fatty acids were extracted twice with 0.5 mL ethyl acetate. The extracted fatty acids were derivatized to fatty acid methyl esters (FAME) by adding 10 μL concentrated HCl, 90 μL methanol and 100 μL of TMS-diazomethane, and incubated at room temperature for 15 min. Gas chromatography-mass spectrometry (GC-MS) analysis of FAME was performed on Agilent 5975 system (Agilent, USA) equipped with a capillary column (DB5-MS, 30 m X 0.25 mm). Sample solutions were analyzed directly by GC-MS at a flow rate of 0.8ml min^-1^, column was equilibrated at 75°C for 1 min, with a 30°C min^-1^ increase to 170°C, 10°C min^-1^ increase to 280°C for holding 2 min. Final FAME concentration was analyzed on the basis of the FAME standard curve obtained from standard FAME mix (GLC-20 and GLC-30, Sigma Aldrich).

### Culture media, cultivation conditions and sample preparation

*P. putida* KT2440 (ATCC 47054) was obtained from ATCC and grown in a chemically defined mineral medium containing the following (per liter): (NH_4_)_2_SO_4_ 1.0 g/L, KH_2_PO_4_ 1.5 g/L, Na_2_HPO_4_ 3.54 g/L, MgSO_4_·7H_2_O 0.2 g/L, CaCl_2_·2H_2_O 0.01 g/L, ammonium ferric citrate 0.06 g/L and trace elements (H_3_BO_3_ 0.3 mg/L, CoCl_2_·6H_2_O 0.2 mg/L, ZnSO_4_·7H_2_O 0.1 mg/L, MnCl_2_·4H_2_O 0.03 mg/L, NaMoO_4_·2H_2_O 0.03 mg/L, NiCl_2_·6H_2_O 0.02 mg/L, CuSO_4_·5H_2_O 0.01 mg/L (56, 57). For biological assays, the BCD fractions was used at concentrations of 80% (v/v) in 15-mL culture tubes. For comparison, additional assays were conducted with (1) *p*-coumarate (1.4 g/L); (2) amino acid mixtures (alanine, serine, proline, glutamic acid, isoleucine, phenylalanine, tyrosine, histidine, tryptophan, threonine, leucine and lysine, total concentration was 0.3 g/L); and (3) the mixture *p*-coumarate (1.4 g/L)/amino acids (0.3 g/L), which were representative of the concentration of these constituents of the BCD fraction diluted to 80% (v/v). Each culture media was sterilized by filtration through 0.20 μm pore size (Thermo Fisher Scientific). Seed cultures for *P. putida* KT2440 were prepared in LB (lysogeny broth) medium at 30 °C, agitated at 200 rpm overnight. The seed culture (2% v/v) was inoculated into each test tube with 10 mL minimal medium (pH was adjusted to 7.0) to start the cultivations at 30 °C and agitated at 200 rpm.

Samples from the cultivations were collected and centrifuged at defined intervals, and the supernatants were stored at −80 °C for GPC and LC−MS analysis. Biomass concentration was measured as optical density at 600 nm in a SpectraMax M2 spectrophotometer (Molecular Devices, San Jose, CA) using 96-well Costar assay plates (Corning Inc., Corning, NY). Three biological replicates were performed, and the obtained values were corrected with uninoculated controls. For proteome analysis, three biological replicates of each different substrate-grown cells were harvested in mid-log phase.

### Generation of ΔPP3358 deletion strains in KT2440

Hydroxycinnamoyl-CoA hydratase-lyase (*ech*; PP_3358) gene deletion mutants in *P. putida* were constructed by homologous recombination and *sacB* counterselection using the allelic exchange vector pMQ30 (58). Briefly, homology fragments 1kb up- and downstream of the target gene, including the start and stop codons respectively, were cloned into pMQ30 via Gibson assembly. Plasmids were then transformed via electroporation in *E. coli* S17 and then mated into *P. putida* via conjugation. Transconjugants were selected for on LB Agar plates supplemented with gentamicin 30 mg/mL, and chloramphenicol 30 mg/mL. Transconjugants were then grown overnight on LB media also supplemented with gentamicin 30 mg/mL, and chloramphenicol 30 mg/mL, and then plated on LB Agar with no NaCl supplemented with 10% (w/v) sucrose. Putative deletions were restreaked on LB Agar with no NaCl supplemented with 10% (w/v) sucrose, and then were screened via PCR with primers flanking the target gene to confirm gene deletion (Table S3). Plasmids and primers were designed using Device Editor (59) and j5 software (60), and plasmids were assembled with Gibson assembly (61). The strain (JPUB_013613) is available from the JBEI Public Registry (https://public-registry.jbei.org/; Table 2).

### Generation of ΔPP_0308-0313 and ΔPP_5063-5064 deletion strains in KT2440

First, deletion mutant *P. putida* ΔPP5063_5064 was assembled via the same method as previously described in creation of ΔPP3358 deletion strains in KT2440 above. Homologous recombination and *sacB* counterselection was used to construct the double knockout deletion mutants targeting the genes NAD-dependent betaine aldehyde dehydrogenase (betB; PP5063) and choline dehydrogenase (betA-II; PP5064). Transconjugants were screened via colony PCR with primers flanking the target gene deletion regions (Table S4). The deletion of dimethylglycine dehydrogenase (dgc operon; PP0308-0313) in *P. putida* was further constructed by homologous recombination and *sacB* counterselection using the vector pBF_ΔPP_0308-0313. Homologous fragments of 500 bps flanking the up-and downstream regions of the target genes were amplified through PCR and assembled into the pKD019MobSacB vector with restriction enzyme digestion and T4 ligation. The assembled plasmid was transformed via heat-shock into *E. coli* S17 cells and transformants were selected on LB Agar plates supplemented with 50 mg/mL kanamycin. Positive transformants were grown overnight in LB broth supplemented with 50 mg/mL kanamycin and the plasmid pBF_ΔPP0308-0313 was extracted using the Qiagen Miniprep kit. The extracted pBF_ΔPP0308-0313 plasmid was further transformed into the *P. putida* ΔPP5063_5064 mutant via electroporation. Selection on single and double crossover events were screened by plating on LB Agar supplemented with 50 mg/mL kanamycin and LB Agar with no NaCl supplemented with 25% (w/v) sucrose. Deletion mutants that did not grow when restreaked onto kanamycin supplemented plates were confirmed via colony PCR using primers that flank the deletion region in the genome (Table S4). Plasmids and primers were designed using SnapGene (GSL Biotech; available at snapgene.com). In order to compare the genotype of the *P. putida* ΔPP5063_5064_0308-0313 mutant and the wild type *P. putida* KT2440 in the presence of choline, each strain was grown in M9 medium supplemented with 0.2% (w/v) choline aspartate and monitored over 24 hours using a TECAN F200 microplate reader (TECAN, Switzerland) at 30 °C, agitated at 200 rpm. The Strain (JPUB_013615) is available from the JBEI Public Registry (https://public-registry.jbei.org/; Table 2).

### Standard flow mass spectrometry and LC-MS/MS data analysis

Samples prepared for shotgun proteomic experiments were analyzed by using an Agilent 6550 iFunnel Q-TOF mass spectrometer (Agilent Technologies, Santa Clara, CA) coupled to an Agilent 1290 UHPLC system as described previously (62). Twenty (20) μg of peptides were separated on a Sigma–Aldrich Ascentis Peptides ES-C18 column (2.1 mm × 100 mm, 2.7 μm particle size, operated at 60 °C) at a 0.400 mL/min flow rate and eluted with the following gradient: initial condition was 95% solvent A (0.1% formic acid) and 5% solvent B (99.9% acetonitrile, 0.1% formic acid). Solvent B was increased to 35% over 120 min, and then increased to 50% over 5 min, then up to 90% over 1 min, and held for 7 min at a flow rate of 0.6 mL/min, followed by a ramp back down to 5% B over 1 min where it was held for 6 min to re-equilibrate the column to original conditions. Peptides were introduced to the mass spectrometer from the LC by using a Jet Stream source (Agilent Technologies) operating in positive-ion mode (3500 V). Source parameters employed gas temp (250 °C), drying gas (14 L/min), nebulizer (35 psig), sheath gas temp (250 °C), sheath gas flow (11 L/min), VCap (3500 V), fragmentor (180 V), OCT 1 RF Vpp (750 V). The data were acquired with Agilent MassHunter Workstation Software, LC/MS Data Acquisition B.06.01 operating in Auto MS/MS mode whereby the 20 most intense ions (charge states, 2-5) within 300–1,400 m/z mass range above a threshold of 1,500 counts were selected for MS/MS analysis. MS/MS spectra (100-1700 m/z) were collected with the quadrupole set to “Medium” resolution and were acquired until 45,000 total counts were collected or for a maximum accumulation time of 333 ms. Former parent ions were excluded for 0.1 min following MS/MS acquisition.

The acquired proteomic data were exported as mgf files and searched against the latest *Pseudomonas putida* KT2440 protein database with Mascot search engine version 2.3.02 (Matrix Science). The resulting search results were filtered and analyzed by Scaffold v 4.3.0 (Proteome Software Inc.). The normalized spectra count of identified proteins were exported for relative quantitative analysis and selecting target genes which expression were significantly altered between experimental groups. In addition, clusters of orthologous groups of proteins (COG) annotations of our proteomes were subjected to classification using RPS-BLASAhT program on COG database (https://www.ncbi.nlm.nih.gov/COG/). The mass spectrometry proteomics data have been deposited to the ProteomeXchange Consortium via the PRIDE partner repository (63) with the dataset identifier PXD014285 and 10.6019/PXD014285.

## Authors Contributions

MP conducted the experiments, analyzed the data and drafted the manuscript. YC and CJP performed the shotgun proteomic experiments. JMG advised the experimental design for ILs pretreatments and BCD reaction and lignin analysis. VTB and EB conducted the LC-MS/MS measurements. JDK, MT and BF constructed mutant strains and plasmids. PD, BAS and SWS conceived the project. SWS supervised the research and drafted the manuscript.

